# CryoCycle your grids: Plunge vitrifying and reusing clipped grids to advance cryoEM democratization

**DOI:** 10.1101/2024.01.23.576763

**Authors:** Viacheslav Serbynovskyi, Jing Wang, Eugene YD Chua, Aygul Ishemgulova, Lambertus M. Alink, William C. Budell, Jake D. Johnston, Charlie Dubbeldam, Fabio A. Gonzalez, Sharon Rozovsky, Edward T. Eng, Alex de Marco, Alex J. Noble

## Abstract

CryoEM democratization is hampered by access to costly plunge-freezing supplies. We introduce methods, called CryoCycle, for reliably blotting, vitrifying, and reusing clipped cryoEM grids. We demonstrate that vitreous ice may be produced by plunging clipped grids with purified proteins into liquid ethane and that clipped grids may be reused several times for different protein samples. Furthermore, we demonstrate the vitrification of thin areas of cells prepared on gold-coated, pre-clipped grids.

---

Cryo-electron microscopy (cryoEM) of biological specimens requires sample vitrification prior to imaging to preserve high-resolution details of biomolecular structures (Taylor & Glaeser, 1974; Frank, 2006). Conventional cryoEM grid preparation entails applying ∼3 μL of protein sample to an unclipped EM grid, blotting away excess sample, and plunging the grid into liquid nitrogen (LN2)-cooled liquid ethane (commonly using commercial devices such as the Thermo Fisher Scientific (TFS) Vitrobot or Leica EM GP) (Dubochet & McDowall, 1981; Dobro *et al*., 2010). During the screening step to optimize conditions for high-resolution structure determination, grid and sample conditions are varied, dozens of grids are prepared, and grids are either screened in a side-loading cryo-transmission EM (cryoTEM) or clipped and screened in a cryoTEM with an autoloader (Sawh-Gopal *et al*., 2023). Each new screening condition requires one or more new grids and possibly autogrid rings and c-clips. However, initial and recurring grid preparation costs, along with availability limitations of consumables, restrict the process to sufficiently-funded institutions.

One solution to decrease grid preparation costs would be to reuse grid-autogrid assemblies (herein called ‘clipped grids’ in general or ‘pre-clipped grids’ if clipping occurs before freezing). We focused on developing methods to reuse clipped grids without disassembling them. The primary challenges with reusing clipped grids are 1) sample vitrification and 2) cleaning clipped grids. Regarding challenge (1), researchers in academia and industry have attempted to blot and vitrify pre-clipped grids consistently for over a decade without success and with minimal reporting in the literature (Ravelli *et al*., 2020). Regarding challenge (2), while a reported method exists for cleaning unclipped EM grids that also removes carbon film (Goldie *et al*., 2014), no method has been reported for cleaning unclipped or clipped EM grids that does not remove the film. This research gap stems from the absence of an affordable method for vitrifying clipped cryoEM grids, with the only available option being cost-prohibitive ethane jet vitrification, such as the implementation in the VitroJet (Ravelli *et al*., 2020).

We examined existing blotting methods to address challenge (1) by testing multiple blotting materials in multiple instruments (TFS Vitrobot Mark IV and Leica EM GP2) and found that no combination sufficiently thinned the ice to <100 nm uniformly nor consistently vitrified the ice (**Supplementary Fig. 1**). In these tests, a minority of grid holes contained thin, vitreous ice, and many contained broken and/or poorly hydrated ice. The former issue has been speculated to be due to the autogrid’s heat capacity being too high, thus impeding vitrification (Ravelli *et al*., 2020). For the latter issue, we suspect that the blotting paper fails to uniformly contact the grid, causing locally dry areas (**Supplementary Fig. 1**). This is due to the autogrid extending tens to hundreds of microns above the grid (**Supplementary Fig. 2**), causing blotting paper larger than the clipped grid to not uniformly contact the grid.

Here we demonstrate a reliable method for producing consistently thin, vitreous ice by applying sample to a clipped grid, blotting only the grid, and immediately freezing in liquid ethane (solution to challenge (1)) (**Supplementary Video 1**). Blotting the grid just inside of the autogrid assembly leaves a thin film of sample which may be vitrified by conventional plunge freezing. We introduce a modified pipette tip called a ‘blotting pipette tip’ to blot the grid inside of the clip ring (**Fig. 1a,b; Supplementary Fig. 3**). Subsequently, we demonstrate a method for reusing clipped grids multiple times by washing sequentially with water then isopropanol while shaking (solution to challenge (2)) (**Fig. 2**). We provide protocols for preparing (**Supplementary Protocol 1**) and reusing (**Supplementary Protocol 2**) clipped grids for single particle cryoEM. Furthermore, we show that the blotting method is sufficient for vitrifying thin areas of cells grown on grids and provide a protocol for preparing cell-compatible, pre-clipped grids (**Supplementary Protocol 3**). **Figure 1c-h** shows examples of vitrified samples of single particles (globular proteins and a long complex) and cells (human cells grown on grids; **Supplementary Fig. 4**) prepared on multiple different grid types using blotting pipette tips. We compared orientations of the p97/selenos complex prepared with a blotting pipette tip (**Fig. 1f**) versus conventionally (**Supplementary Fig. 5a**) and found them to be comparable (**Supplementary Fig. 5b,c**). These methods, called CryoCycle, significantly reduce costs and demand for consumables, thus democratizing cryoEM by enabling more widespread and uninterrupted adoption.

**Figure 1.**
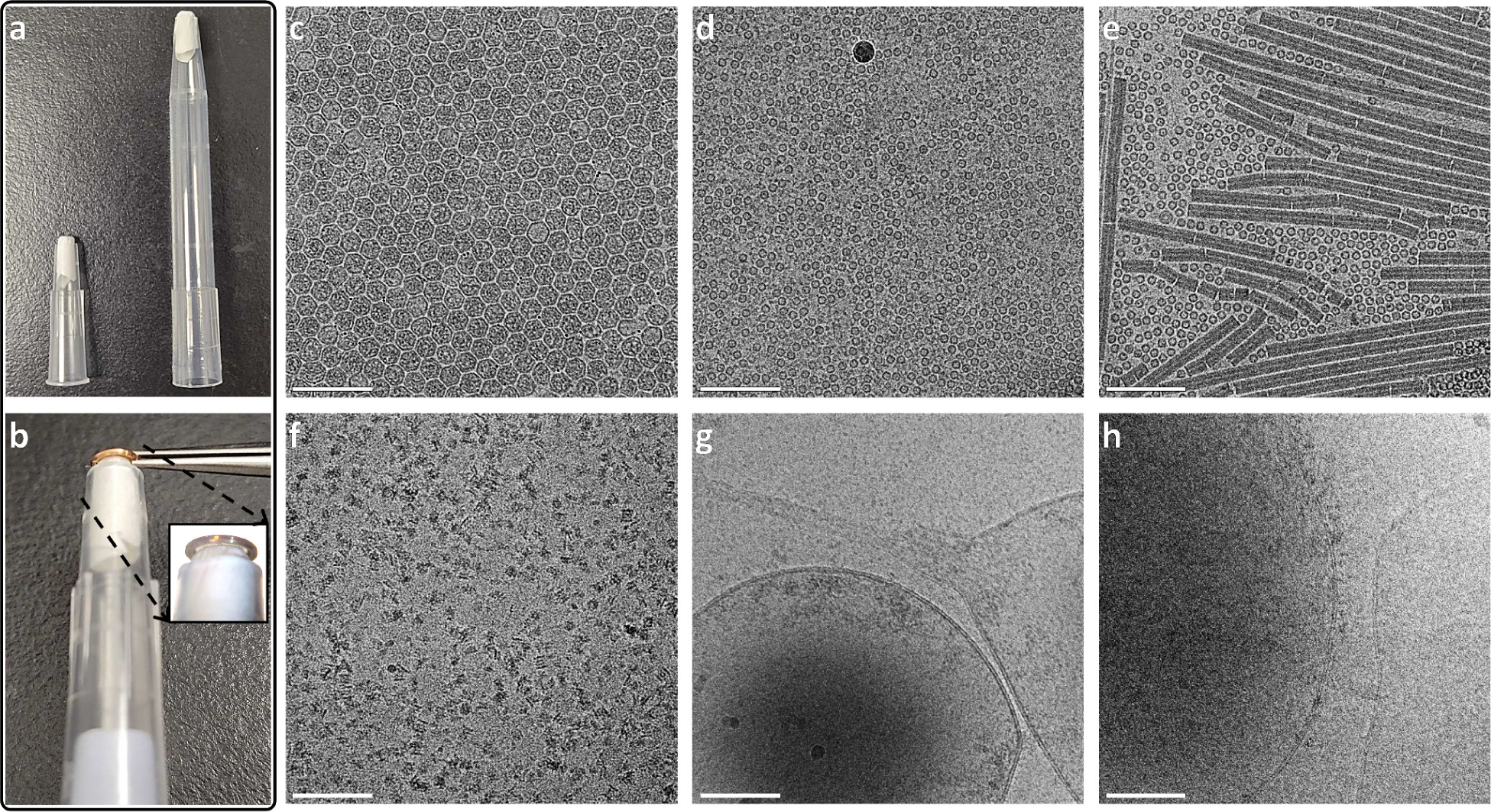
| CryoCycle clipped grid blotting with various designs, grids, and samples. **(a)** Two blotting pipette tip sizes: Trimmed 200 μL (left) and 1,000 μL (right) tips, each with a piece of filter paper inserted for blotting 3+ μL of sample. **(b)** A 200 μL blotting pipette tip applied to the same side of the grid as the single particle sample; see orientations in **Supplementary Figure 3a**. The inset shows the contact between the blotting paper and the grid inside the autogrid ring. *Note*: Blotting for cell samples is from the opposite side of the grid, as shown in **Supplementary Figure 3b**. **(c-h)** A selection of micrographs of vitreous single particle and cell samples on multiple grid types (gold and carbon) prepared with the CryoCycle method; **(c)** Virus-like Particles (VLPs), **(d)** Apoferritin, **(e)** A protein mixture of apoferritin and tobacco mosaic virus (TMV), **(f)** A p97/selenos complex (conventional preparation comparison in **Supplementary Fig. 5**), **(g-h)** Thin edges of human retinal pigment epithelial-1 (RPE-1) cells, showing intact membrane bilayers (cryoFLM images shown in **Supplementary Fig. 4**). Scale bars: 100 nm for (c-h).

**Figure 2.**
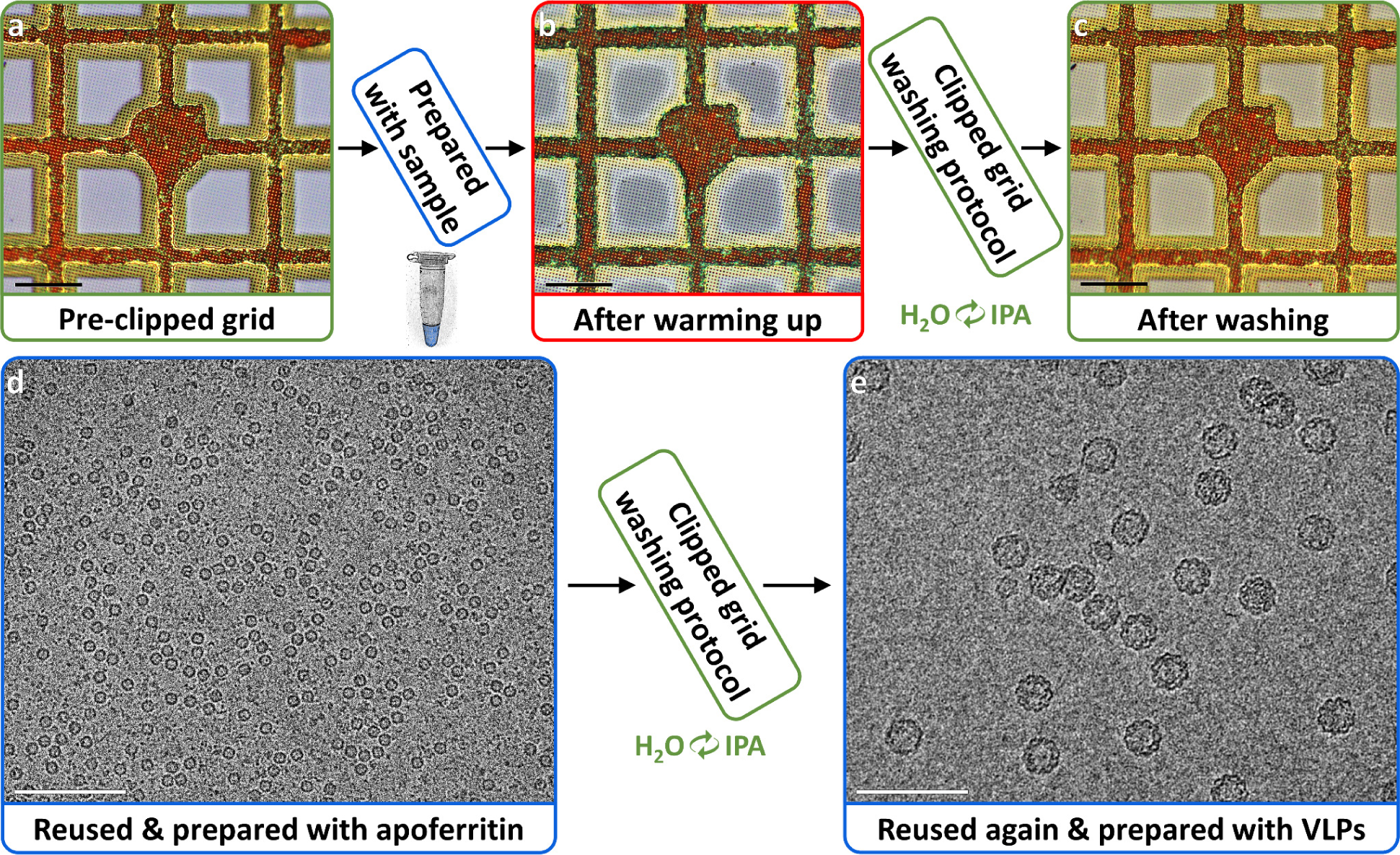
| CryoCycle reused clipped carbon grids washing protocol results. **(a)** Squares of a freshly pre-clipped carbon grid. **(b)** Squares of the same grid after vitrifying a sample with the CryoCycle method, warming up, and drying. **(c)** Squares of the same grid after the washing protocol. **(d)** A micrograph of a clipped grid washed and prepared with apoferritin. **(e)** A micrograph from the clipped grid in (d) washed again and prepared with VLPs, showing VLPs of expected sizes. Scale bars: 50 μm for (a-c), 100 nm for (d-e). **Supplementary Figure 9** shows CryoCycle reused grid washing for a gold grid. **Supplementary Figure 10** shows grid atlases of (d) & (e).

The key components for blotting clipped grids for vitrification are rigid, flat-tipped tweezers and a blotting pipette tip that uniformly contacts the grid. For the former, we found that conventional fine-tipped Vitrobot tweezers insecurely hold clipped grids due to their flexible tips, causing rotation into the tweezers when handled (**Supplementary Fig. 6a,b**). Trimming the tweezer tips increases rigidity sufficiently to handle the autogrid without rotating into the grid, which is essential for blotting (**Supplementary Fig. 6a,c & 7; Supplementary Video 1; Supplementary Protocol 1**). For the latter, a blotting pipette tip is created by cutting the narrow end of a pipette tip to a 3 mm inner diameter then inserting a ∼1 cm piece of filter paper until 1 mm protrudes out of the small end (**Fig. 1a,b; Supplementary Protocol 1**). Rounding the protruding filter paper’s edges facilitates blotting within the clip ring, while flattening the end helps ensure uniform grid contact (**Supplementary Fig. 8**). 200 µL and 1,000 µL blotting pipette tips accommodate enough blotting paper to absorb 3+ µL of sample (**Fig. 1a,b**). **Supplementary Video 2** shows the CryoCycle blotting pipette tip assembly process. **Supplementary Figure 3** depicts sample application, blotting direction, and orientation of the grid and autogrid for single particle and cell samples.

We developed a washing protocol for reusing single particle clipped grids, taking into account the sensitivity of the grid film and the autogrid assembly. Clipped EM grids are typically composed of copper, carbon, gold, and palladium. Washing them first with water removes bulk contamination, then subsequent isopropanol washes displace and rinse away remaining samples and contaminants. Isopropanol was selected for its minimal reactivity with these metals at room temperature and high purity (99+%) to prevent altering their chemical composition. We determined that one 5-minute wash with water followed by two 5-minute isopropanol washes sufficiently cleans clipped carbon and gold grids (**Fig. 2; Supplementary Fig. 9; Supplementary Protocol 2**). **Figure 2e** shows a micrograph from a clipped carbon grid that was reused twice (i.e. 3 samples, 2 washings) and **Supplementary Figure 10** shows their grid atlases with the vast majority of squares intact, exemplifying the protocol’s robustness. The maximum number of times clipped grids may be reused has not yet been determined.

The CryoCycle clipped grid preparation methods offer significant advantages over conventional unclipped grid preparation in four key areas. 1) *Room temperature grid clipping is simpler and safer*, as it avoids mechanical and visual disruptions caused by LN2, enables easy visual verification of clipped ring placement by eye, and eliminates risks of grids thawing, fingers freezing, and ice contamination. Additionally, a clean, flat surface can be used to clip instead of a clipping station, saving a one-time cost of thousands of dollars. 2) *Handling pre-clipped grids mitigates mechanical damage and user stress* (**Supplementary Fig. 11**) compared to handling unclipped grids (**Supplementary Fig. 7**); ideally, a grid is handled once for clipping, then only the autogrid is touched thereafter. Handling clipped grids substantially reduces bent grids, which expedites screening due to minimal defocus gradients. Moreover, clipped grid handling eliminates the need to purchase unclipped grid boxes. 3) *CryoCycle blotting and freezing only requires a humidity chamber*; popular commercial semi-automated plunge freezing devices are not needed, which may reduce one-time costs by tens of thousands of dollars. While we illustrate the use of CryoCycle in a Vitrobot, we emphasize that only the humidity chamber and plunger are used (**Supplementary Video 1**); CryoCycle grid preparation using a gravity plunger (Comolli *et al*., 2012; Depelteau *et al*., 2020) or manually plunging by hand is possible, although the latter has been minimally tested (**Supplementary Fig. 12**). 4) *Reusing clipped grids reduces initial and recurring costs*. Each pre-clipped grid assembly costs about $45 in 2024, so for a typical cryoEM sample where 24 grids are required to be screened before conditions suitable for data collection are found, reusing 8 grids three times each would reduce recurring costs by $720. Most cryoEM projects require optimization of several samples, thus using CryoCycle methods reduces costs of cryoEM projects by thousands of dollars per project and reduces costs for cryoEM labs by tens of thousands of dollars per year. **Table 1** lists where CryoCycle methods can reduce both initial and yearly costs by tens of thousands of dollars each for typical cryoEM labs. Additionally, CryoCycle methods enable vitrification of clipped grids by plunge freezing, circumventing the previous requirement of using ethane jet freezers that cost hundreds of thousands of dollars. The advantages listed here particularly benefit new cryoEM users, making cryoEM more accessible.

**Table 1.** | CryoCycle savings versus conventional preparation costs. CryoCycle methods save tens to hundreds of thousands of dollars initially and tens of thousands of dollars yearly for a typical cryoEM lab (bold). Added costs are ∼$5,000 initially (bold), $0 recurring. Grid savings assume reusing clipped grids 3 times each. Conventional preparation may incur additional losses of unclipped grids, varying with user expertise. Staffing costs or savings may vary depending on how the protocols are implemented.

| Potential CryoCycle cost savings |  |  |  |  |  |
| --- | --- | --- | --- | --- | --- |
| Item | One-time cost saved | Recurring cost (per item) | Recurring cost (per year) | Recurring cost saved (per year) | Notes |
| Commercial semi-automated plunge freezer | **~\$100,000** | -- | -- | -- | Approx. Leica EM GP or Vitrobot cost. Common in facilities. |
| Jet freezer | >\$500,000 | -- | -- | -- | Approx. VitroJet cost. For comparison of an existing method to vitrify clipped grids. |
| Clipping station | **\$2,500** | -- | -- | -- | Approx. cost of a new clipping station. |
| CryoEM grids | -- | \$10-20 | \$6,000 | **\$4,000** | Assuming 400 grids per year (~8 grids per week) and an avg. cost of \$15 per grid. |
| Clip rings and c-clips | -- | \$30 per pair | \$12,000 | **\$8,000** | Assuming 400 rings and c-clips per year. |
| Unclipped grid boxes | -- | \$10 | \$100 | **\$100** | Assuming reuse of 10 grid boxes per year. |
| Potential additional CryoCycle costs |  |  |  |  |  |
| Item | One-time cost | Recurring cost (per item) | Recurring cost (per year) | Recurring cost saved (per year) | Notes |
| Rotary tool and disk | **\$200** | -- | -- | -- | Common in machine shops. |
| Tweezer sharpening tool | **\$100** | -- | -- | -- | Common in cryoEM labs. |
| CryoCycle tweezers | **\$700** | -- | -- | -- | Described in Supplementary Protocol 1 |
| Plunge freezer w/ humidity control | **\$4,000** | -- | -- | -- | E.g. Comolli *et al.*, 2012; Depelteau *et al.*, 2020. |

## METHODS

### Sample Preparation

#### Virus-like Particles (VLPs)

PP7 Leviviridae PP7 VLPs were prepared as described in (Zhao *et al*., 2019) at ∼1 mg/mL.

#### Apoferritin

Mouse apoferritin stock (8 mg/mL) was provided by Dr. Kikkawa’s lab (U Tokyo).

#### Protein mixture

Purified apoferritin and tobacco mosaic virus (TMV) were mixed in phosphate buffered saline at concentrations of 2.9 mg/mL and 13.6 mg/mL, respectively.

#### p97/selenos complex

The AAA ATPase p97 was combined with an excess molar quantity of full-length selenoprotein S U188C (selenos), resulting in a final concentration of the p97/selenos complex at 11 mg/mL. This mixture was incubated on ice for 30 minutes, including 5 mM ATP*γ*S and 1.2 mM DDM (1 CMC), prior to grid application. The preparation of selenos followed the method outlined in (Ghelichkhani *et al*., 2023).

#### Human retinal pigment epithelial-1 (RPE-1) cells

RPE-1 cells expressing IRFP::hCentrin2, mCherry::α-tubulin, sfGFP::CENP-A were acquired from the Needleman Lab (Harvard University) and cultured in DMEM (Thermo Fisher Scientific) supplemented with 10% FBS (Thermo Fisher Scientific) and Penicillin-Streptomycin (Thermo Fisher Scientific) in a humidified incubator with 5% CO_2_ maintained at 37°C. Cells were trypsinized and plated onto gold-coated, pre-clipped carbon Quantifoil grids (**Supplementary Protocol 3**) at a density of 1,000 cells per grid and grown overnight prior to freezing.

### Grid Preparation and Vitrification

For samples prepared with the CryoCycle blotting method inside a TFS Vitrobot Mark IV chamber (Thermo Fisher Scientific) (**Fig. 1c-h, Fig. 2d,e, Supplementary Fig. 4**), all time and force parameters were set to zero (Blot Time, Blot Force, Wait Time, and Drain Time) and humidity was set to 80%. The instrument was operated at room temperature. Quantifoil (Quantifoil Micro Tools GmbH) holey carbon, gold, and UltrAuFoil grids were used. Single particle samples were blotted on the same side as sample application following the Procedure in **Supplementary Protocol 1** and cell samples were blotted on the opposite side as cell growth (see **Supplementary Fig. 3**).

For samples prepared conventionally using multiple blotting materials in the TFS Vitrobot Mark IV and Leica EM GP2 chambers (**Supplementary Fig. 1**), several parameters were varied across different test samples (primarily apoferritin, VLPs, and the protein mixture) including: Quantifoil and C-Flat (Protochips Inc.) carbon and gold grids, blot times from 1 to 4 seconds, and 70-85% humidity.

### CryoEM Grid Screening

Samples were screened on a TFS Glacios cryoTEM (Thermo Fisher Scientific) and a TFS Falcon 3EC camera operating in integrating mode (for most samples) or counting mode using either Smart Leginon Autoscreen (Cheng *et al*., 2023) or manually in Leginon (Potter *et al*., 1999). Collection parameters varied across screening sessions; nominal defocus: ∼-1.5 to −3 μm, pixelsize: 1.204 Å/pixel, total dose: 50-60 e-/Å^2^.

### CryoEM Processing

The p97/selenos complex prepared with the CryoCycle method (one UltrAuFoil grid and one Quantifoil carbon grid) and screened on a TFS Glacios cryoTEM resulted in 592 micrographs (integrating mode; no frame alignment). Micrographs were processed in Cryosparc v4.4.1 (Punjani *et al*., 2017) by first CTF correcting, then initial picking with templates and Topaz (Bepler *et al*., 2019), then iterative 2D classification, manual cleanup, and Topaz retraining, before converging to 24,584 particles after 2D classification (**Supplementary Fig. 5c**).

The p97/selenos complex prepared conventionally with a TFS Vitrobot Mark IV (one Quantifoil carbon grid) and screened on a TFS Glacios cryoTEM resulted in 2,392 micrographs (counting mode). Micrographs were processed in Cryosparc v4.4.1 (Punjani *et al*., 2017) before selecting a random subset of 24,584 particles for a final 2D classification (**Supplementary Fig. 5b**) to match the number of particles from the CryoCycle method dataset.

### SEM imaging

SEM images (**Supplementary Fig. 1**, 3rd row, 1st column; **Supplementary Fig. 2**) were collected on an EMCrafts Cube II (EmCrafts Co. Ltd).

### CryoFLM imaging

CryoFLM images (**Supplementary Fig. 4**) were collected on a Zeiss LSM 900 with Airyscan 2 (Carl Zeiss Microscopy GmbH) configured to excite the sfGFP bound to the CENP-A nucleosomes.

### Verification of Consent

Photo (**Supplementary Fig. 12a**) and **Supplementary Videos 1 & 2** of Viacheslav Serbynovskyi were taken with approval and are shown here with consent.

## ACKNOWLEDGEMENTS

The authors wish to thank Dr. Bridget Carragher (Chan Zuckerberg Imaging Institute) for early advice in the project, Gloria Ha and Dr. Daniel J. Needleman (Harvard University) for kindly providing the RPE-1 cells, Dr. William Conway (Flatiron Institute; NYSBC) for helping prepare the RPE-1 cells and for critical reading of the manuscript, and the entire SEMC team for helpful discussions. F.A.G. and S.R. were supported by a grant from the NIH National Institute of General Medical Sciences (1R01GM121607-01) and NSF (2150863). Some of this work was performed at the Simons Electron Microscopy Center at the New York Structural Biology Center, with support from the Simons Foundation (SF349247), NIH (U24 GM129539), and NY State Assembly.

## AUTHOR CONTRIBUTIONS

V.S. conceived of the project. A.I. and J.J. prepared the protein mixture sample. F.A.G. and S.R. prepared the AAA ATPase sample. J.J. grew the cells on the grids. V.S. F.A.G., and A.J.N. prepared cryoEM grids. A.I. imaged clipped grids in a LM. C.D. imaged cell cryoEM grids in a cryoFLM. V.S., J.W., E.Y.D.C., and A.J.N. imaged grids in a cryoTEM. L.M.A., W.C.B., E.T.E., A.D.M, and A.J.N. provided oversight. A.J.N. wrote the manuscript and all authors edited the manuscript.

## SUPPLEMENTARY INFORMATION

**Supplementary Figure 1.**
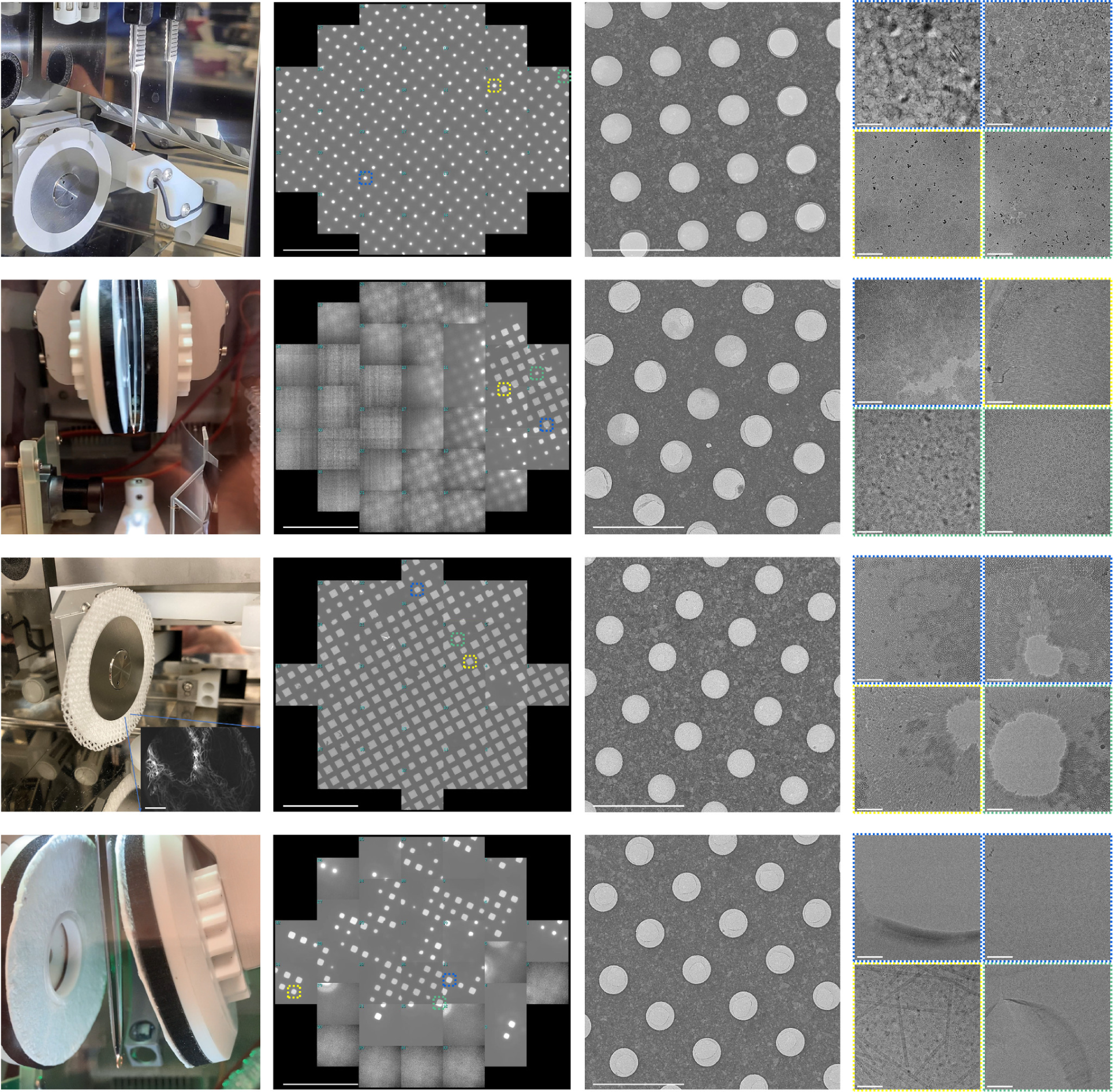
| Pre-clipped grid vitrification attempts with conventional plunge freezers and different blotting papers. First column shows blotting papers tested. Second column shows representative grid atlases with high-magnification areas outlined. Third column shows a representative medium-mag image of grid holes. Fourth column shows representative high-mag images of ice in holes. First row shows normal Leica EM GP2 blotting paper. Second row shows normal Vitrobot blotting paper. Third row shows custom holey cotton paper on top of normal Leica EM GP2 blotting paper. Fourth row shows thick cotton blotting paper. Each attempt resulted in predominantly non-vitreous ice and frequent absence of ice at the centers of holes (3rd column). Scale bars: 500 µm, 100 µm, and 100 nm, respectively for 2nd-4th columns; 500 µm for inset SEM image in 3rd row, 1st column.

**Supplementary Figure 2.**
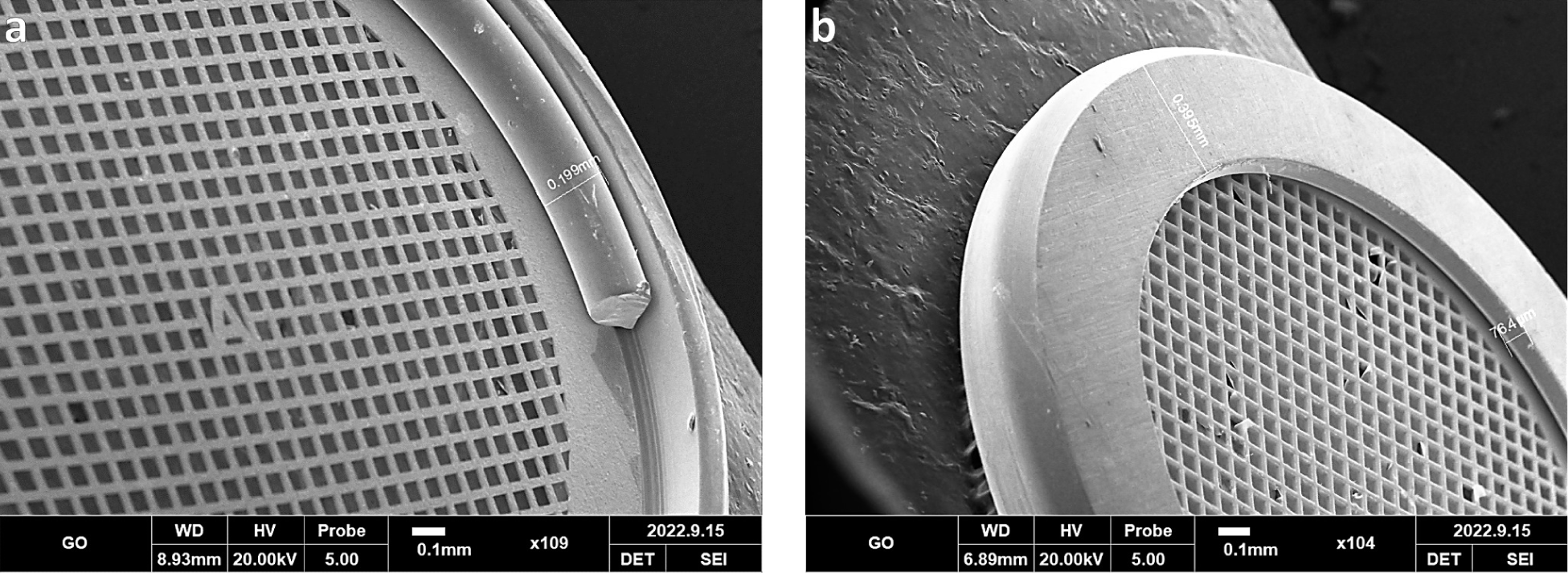
| Measurements from angled SEM images of a clipped grid. **(a)** The front side of the clipped grid shows that the autogrid assembly extends over 100 microns above the grid. **(b)** The back side of the clipped grid shows that the autogrid ring extends many tens of microns above the grid.

**Supplementary Figure 3.**
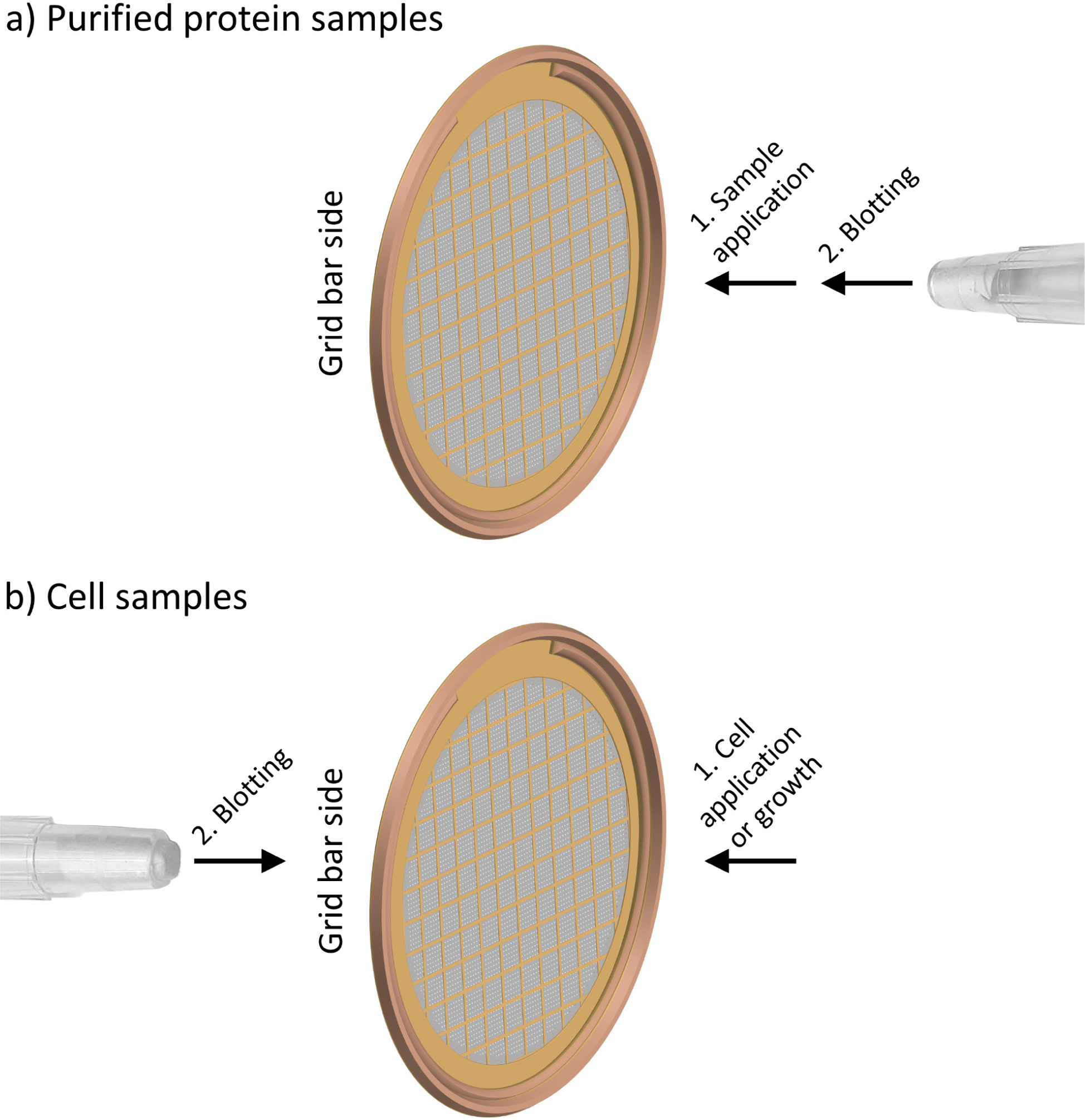
| Grid, autogrid ring and c-clip, sample application, and blotting orientation guide. For both purified protein and cell samples, the grid is clipped such that the grid bars are facing the opposite direction as the c-clip. Sample is applied or adherent cells are grown on the non-grid bar side of the grid. The only difference is that purified protein samples are blotted on the same side as sample application **(a)** and cell samples are blotted from the opposite side as cell application/growth **(b)**. Note: The blotting pipette tips and the clipped grids are not to scale.

**Supplementary Figure 4.**
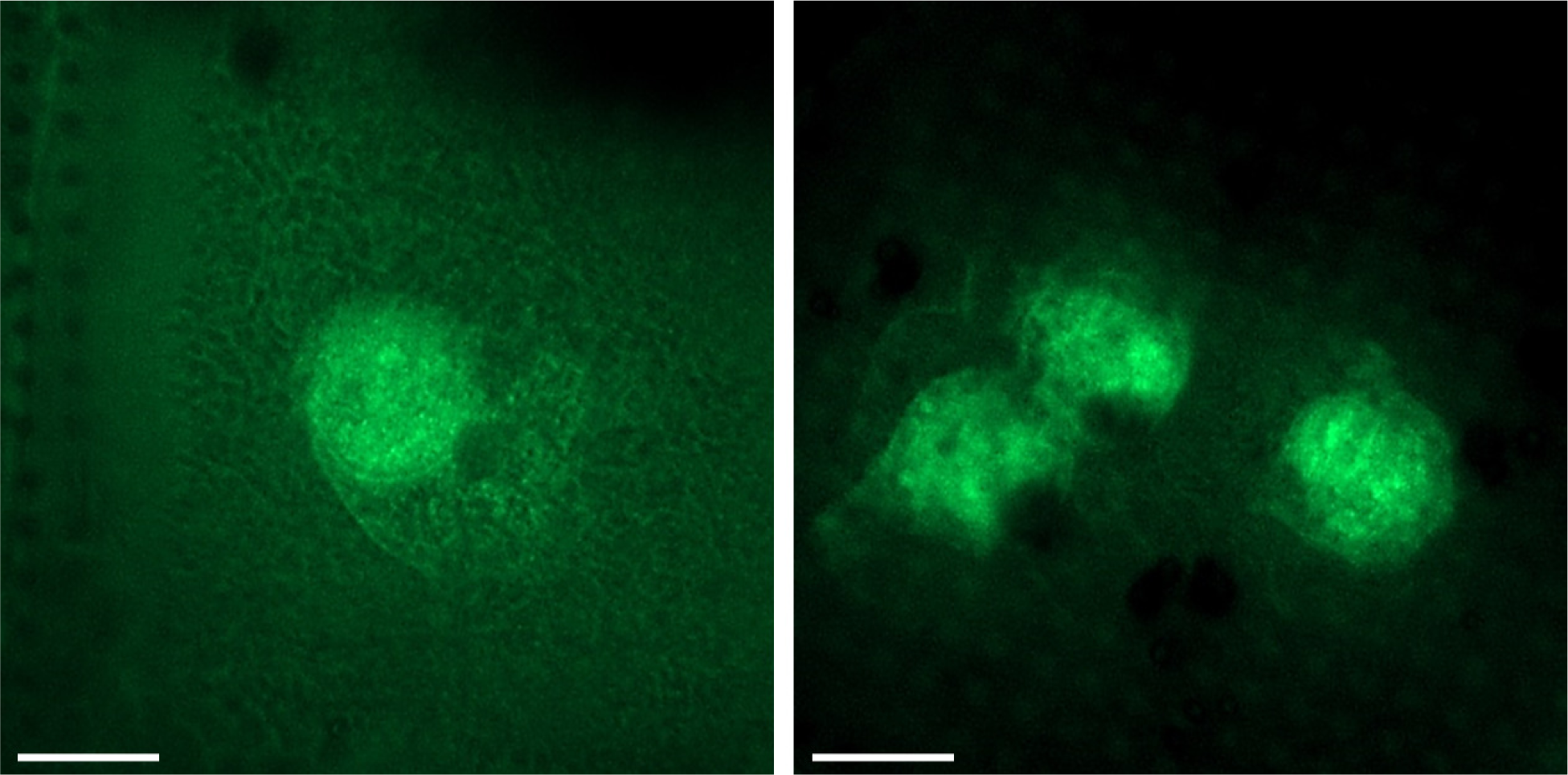
| Cryo-fluorescent light microscopy (cryoFLM) images of cells grown on pre-clipped, gold-coated grids. Two cryoFLM images of RPE-1 cells with CENP-A nucleosomes fluorescing in green from the same grid as shown in **Figure 1g,h**. Delineated organelles can be seen. Scale bars are 10 µm.

**Supplementary Figure 5.**
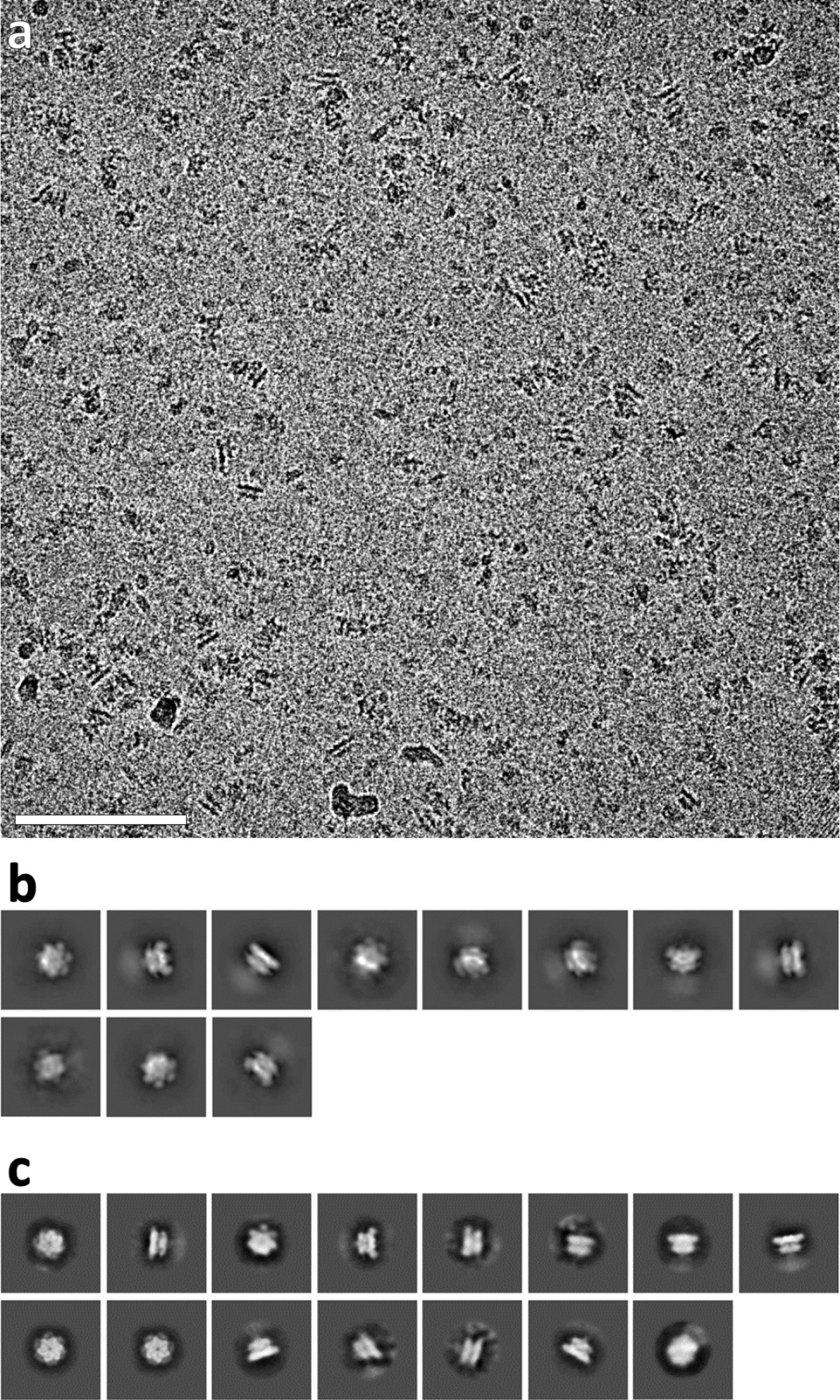
| Comparison of conventional plunge freezing of a p97/selenos complex with plunge freezing a preclipped grid with a blotting pipette tip. **(a)** An example micrograph of the same p97/selenos complex sample in **Figure 1f**, except conventionally prepared with a Vitrobot. **(b)** 2D classes from the dataset in (a). **(c)** 2D classes from the dataset in **Figure 1f**, prepared with a blotting pipette tip. Scale bar: 100 nm.

**Supplementary Figure 6.**
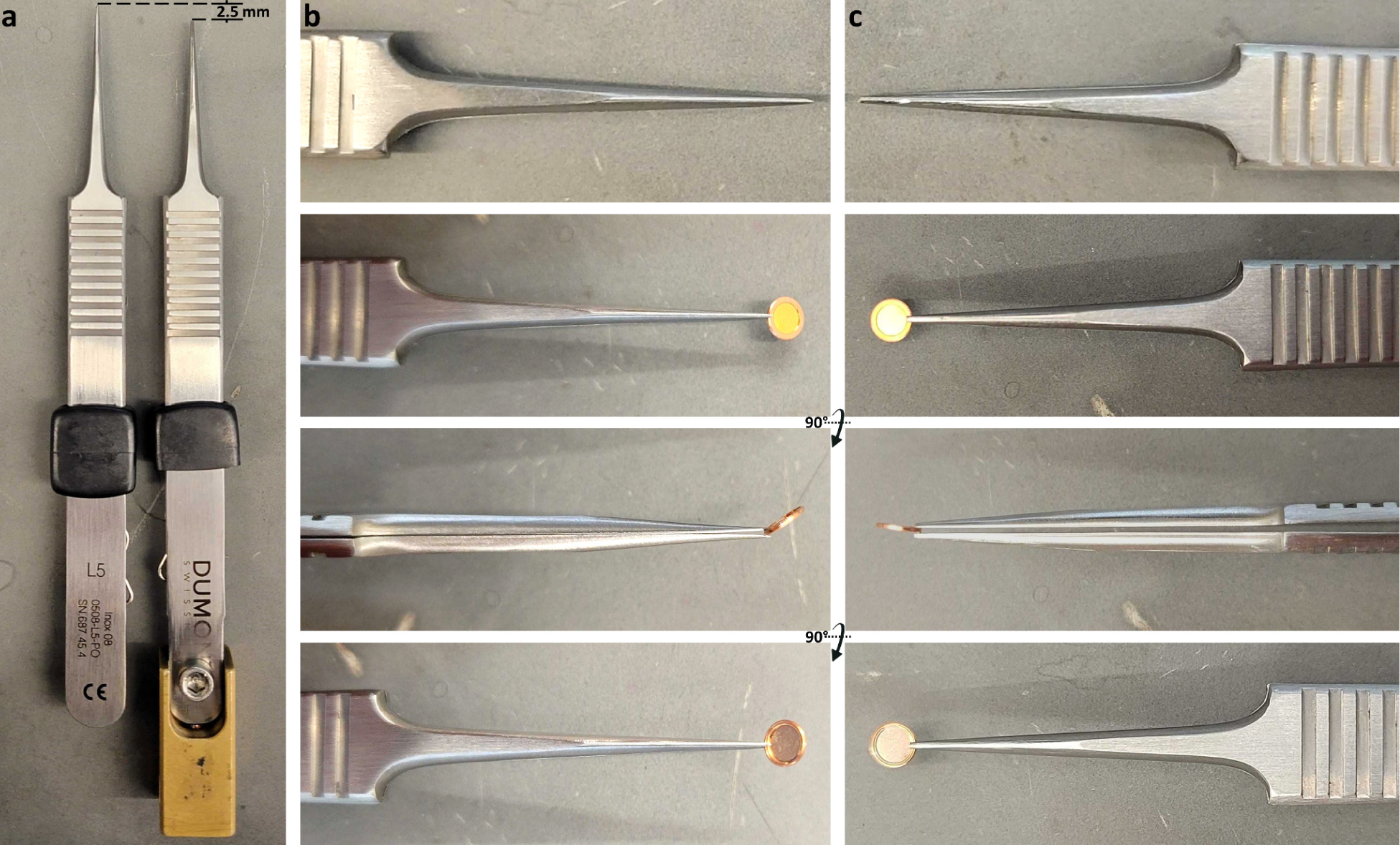
| Comparison of unmodified tweezers and modified tweezers for clipped grid freezing. **(a)** Left: Unmodified Dumont L5 tweezers. Right: Dumont L5 tweezers attached to a Vitrobot mount and modified by trimming off ∼2.5 mm of the tip of the tweezer (**Supplementary Protocol 1**). **(b)** Unmodified tweezers; first row shows the fine tip, second row shows the tweezers holding an autogrid from the point of view of the flat side of the ring, third row is rotated 90° from the second row, fourth row is rotated 90° from the third row. **(c)** Modified tweezers; first row shows the trimmed, flattened tip, second row shows the tweezers holding an autogrid from the point of view of the flat side of the ring, third row is rotated 90° from the second row, fourth row is rotated 90° from the third row. In the third row, the unmodified tweezers bend near the tip (b), causing a ∼45° rotation of the autogrid which prevents easy blotting and risks the tweezers damaging the grid, while the modified tweezers do not bend and the flattened tip securely holds the autogrid with nearly no rotation (c), allowing for CryoCycle blotting.

**Supplementary Figure 7.**
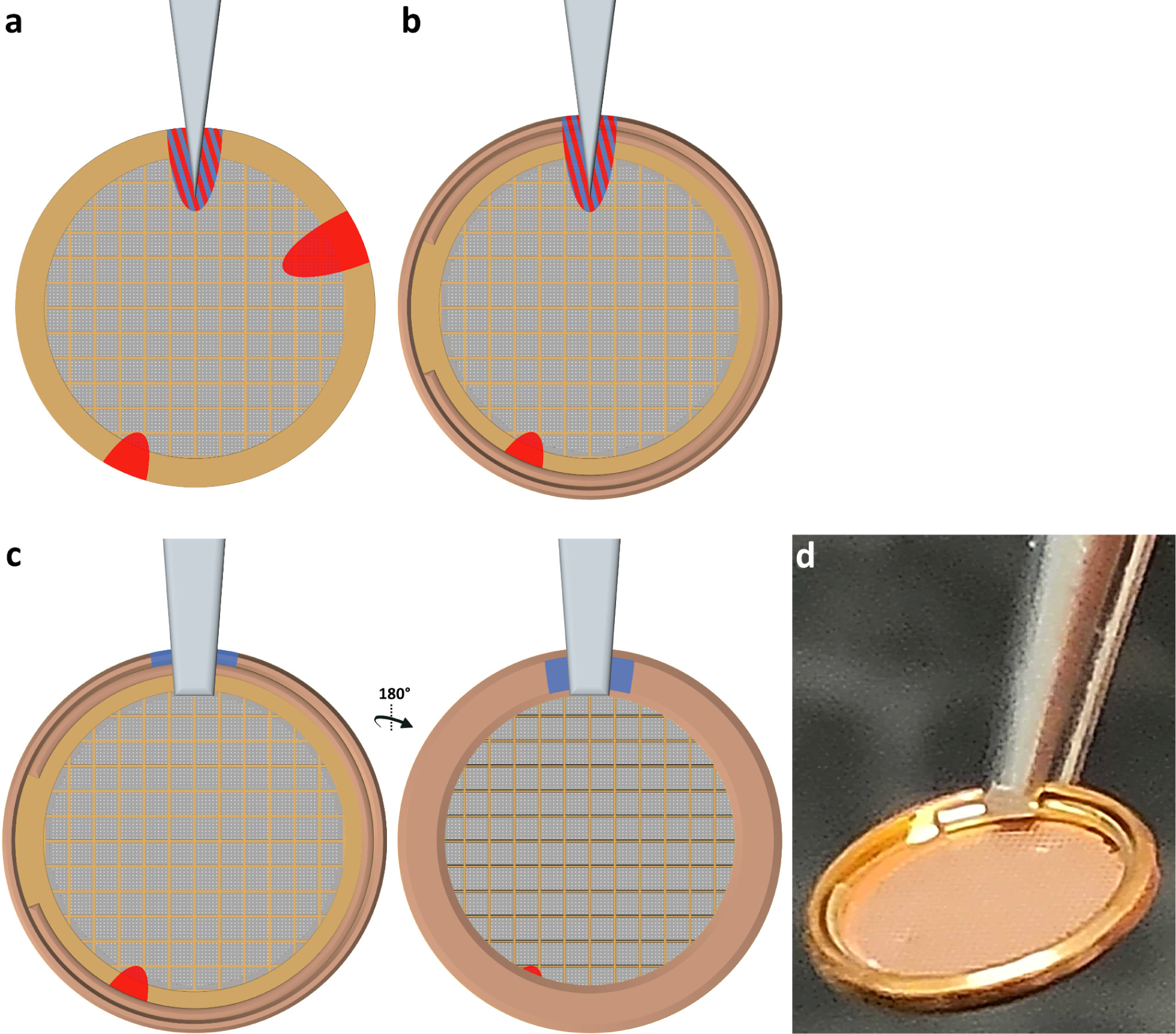
| Estimated areas directly affected mechanically and thermally by tweezer handling. **(a)** A typical grip on an EM grid by fine-tipped tweezers. During several handling steps before imaging (initial grid retrieval, after plasma cleaning/glow discharging to plunge freeze and transfer to a grid box, and handling for clipping), the grid is handled in at least three locations around the rim, potentially causing local physical damage (red). The bottom-left red damage area is from handling at room temperature when it is easier to localize the rim, thus the area is smaller. During freezing, squares near the tweezers are usually not vitrified (red & blue). **(b)** A pre-clipped grid held by fine-tipped tweezers is at high risk of damage near the tweezers, as shown in **Supplementary Figure 6b**, and the grip itself is unstable. **(c)** A clipped grid held by trimmed tweezers (**Supplementary Fig. 6c & Supplementary Protocol 1**) has minimal risk of mechanical damage and minimal risk of vitrification issues (blue) from the tweezers. Both sides of the clipped grid are shown. **(d)** An oblique view of a clipped grid handled by trimmed tweezers as in (c).

**Supplementary Figure 8.**
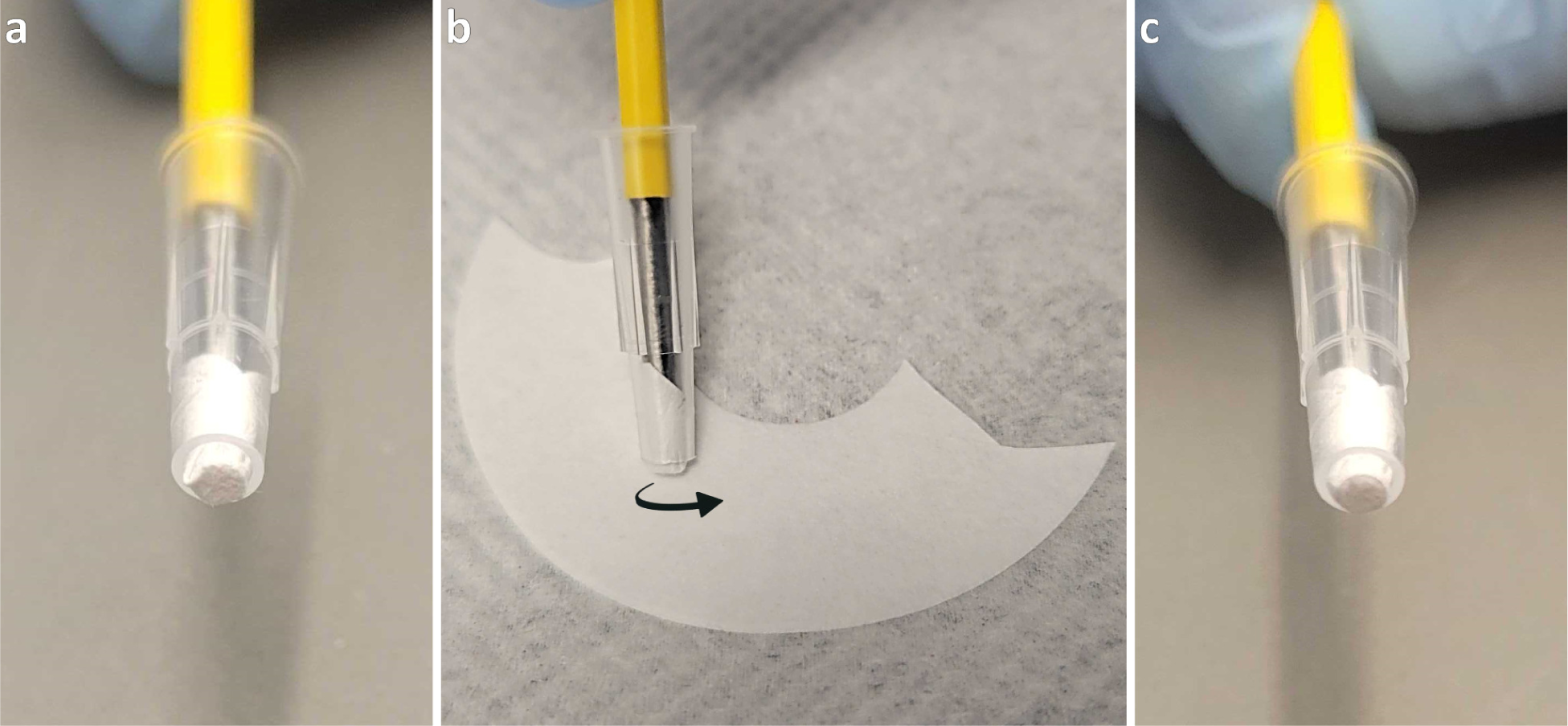
| Rounding the edges of blotting paper in a modified pipette tip. During preparation of the blotting pipette tip (**Supplementary Protocol 1**), it is important to ensure that the edges of the blotting paper extruding from the modified pipette tip are rounded and smoothed to facilitate blotting inside of the clip ring (**Fig. 1b**). To do this, position the assembly at ∼60° with respect to a flat surface with the metal rod inserted and roll the end of the blotting paper on clean filter paper until smooth. **(a)** 200 µL blotting pipette tip before rolling. **(b)** Rolling the blotting tip at a 60° angle on a flat, clean surface. **(c)** Blotting pipette tip after rolling at 60° and pressing at 90°.

**Supplementary Figure 9.**
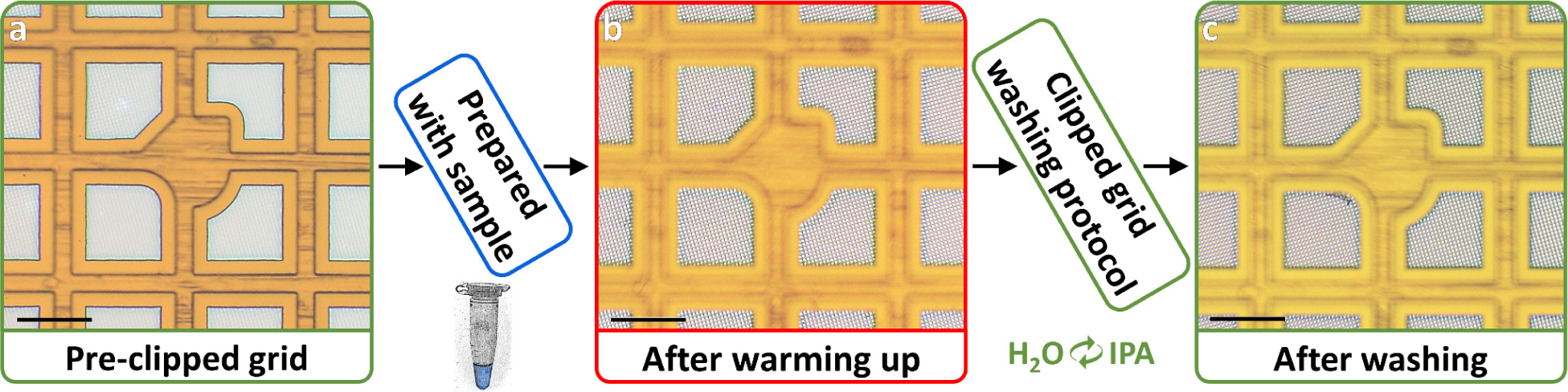
| CryoCycle reused clipped gold grids washing protocol results. **(a)** Squares of a freshly pre-clipped gold grid. **(b)** Squares of the same grid after vitrifying a sample with the CryoCycle method, warming up, and drying. **(c)** Squares of the same grid after the washing protocol.

**Supplementary Figure 10.**
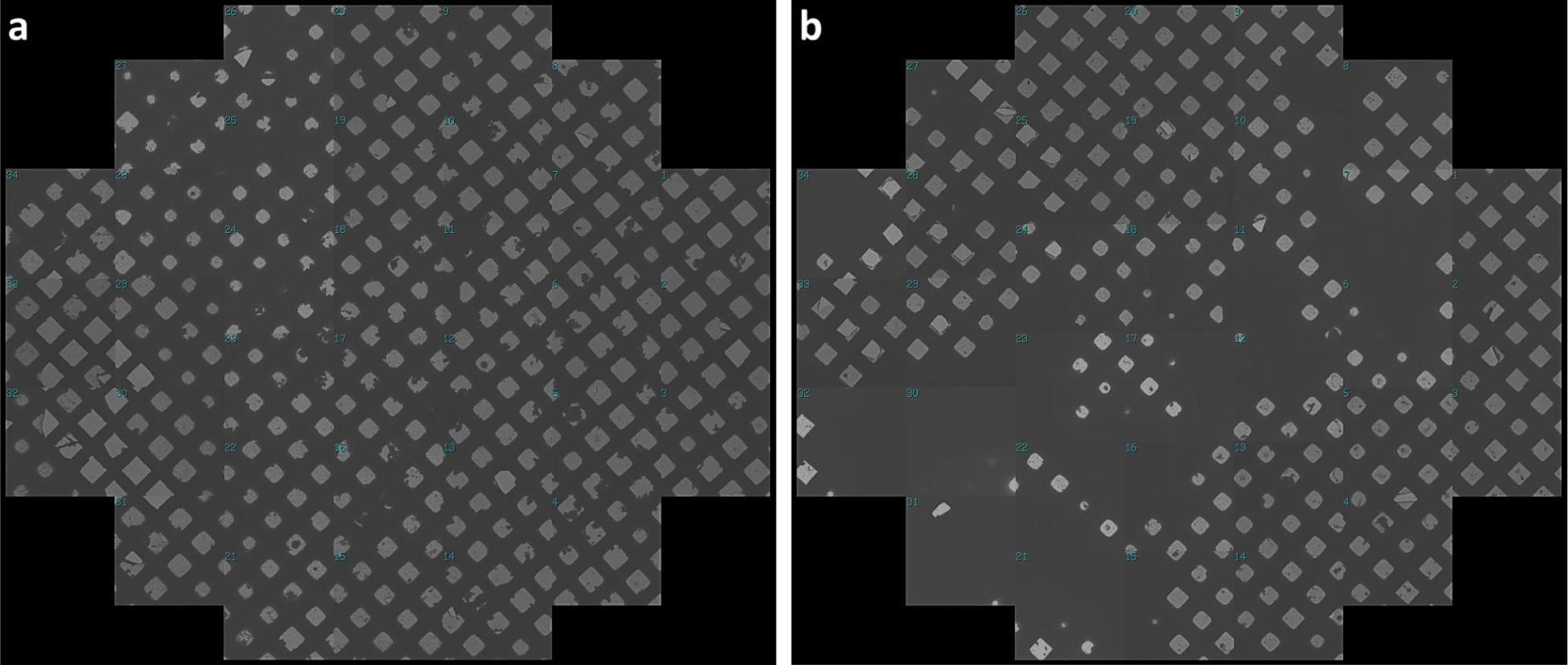
| Grid atlases of a carbon grid after washings. **(a)** Grid atlas of the carbon grid with apoferritin shown in **Figure 2d** that has already been washed and reused once - after preparation with VLPs - showing that the vast majority of grid squares are intact. **(b)** Grid atlas of the same grid shown in **Figure 2e** that was washed and reused again and prepared with VLPs showing a similar small number of broken grid squares as in (a). The previous sample was not found to be on the grid after the washings.

**Supplementary Figure 11.**
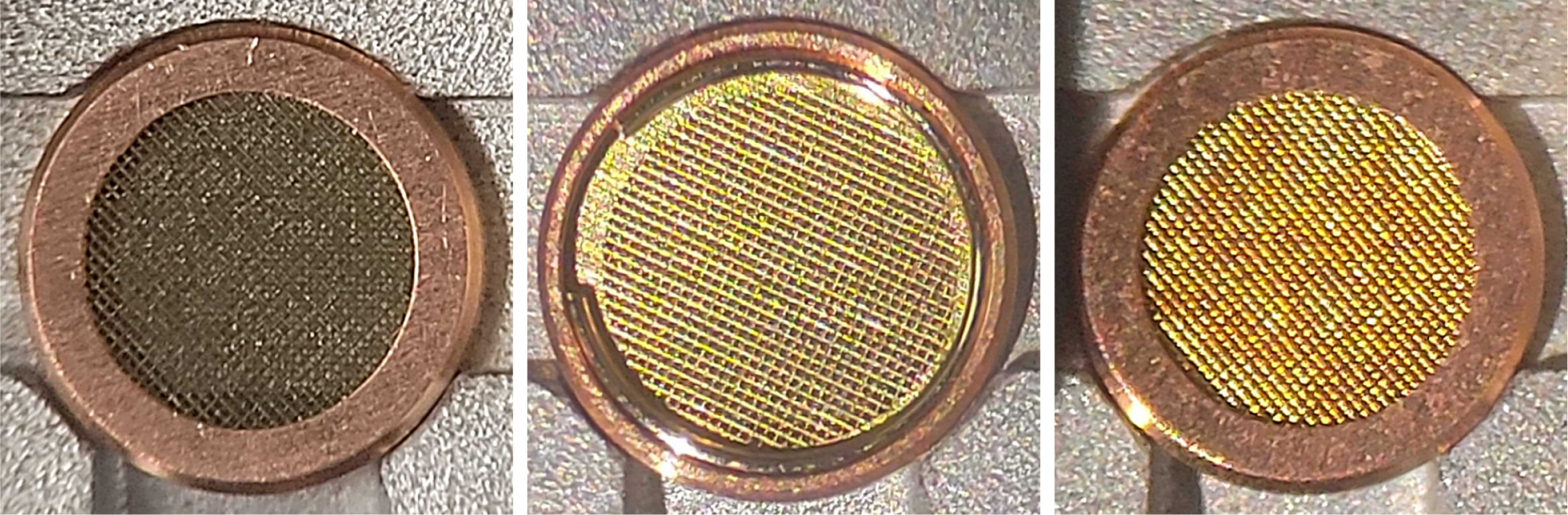
| Examples of grids clipped at room temperature. Photos of one carbon (left) and two gold grids (middle, right) clipped at room temperature. The grids show virtually no damage due to minimal handling.

**Supplementary Figure 12.**
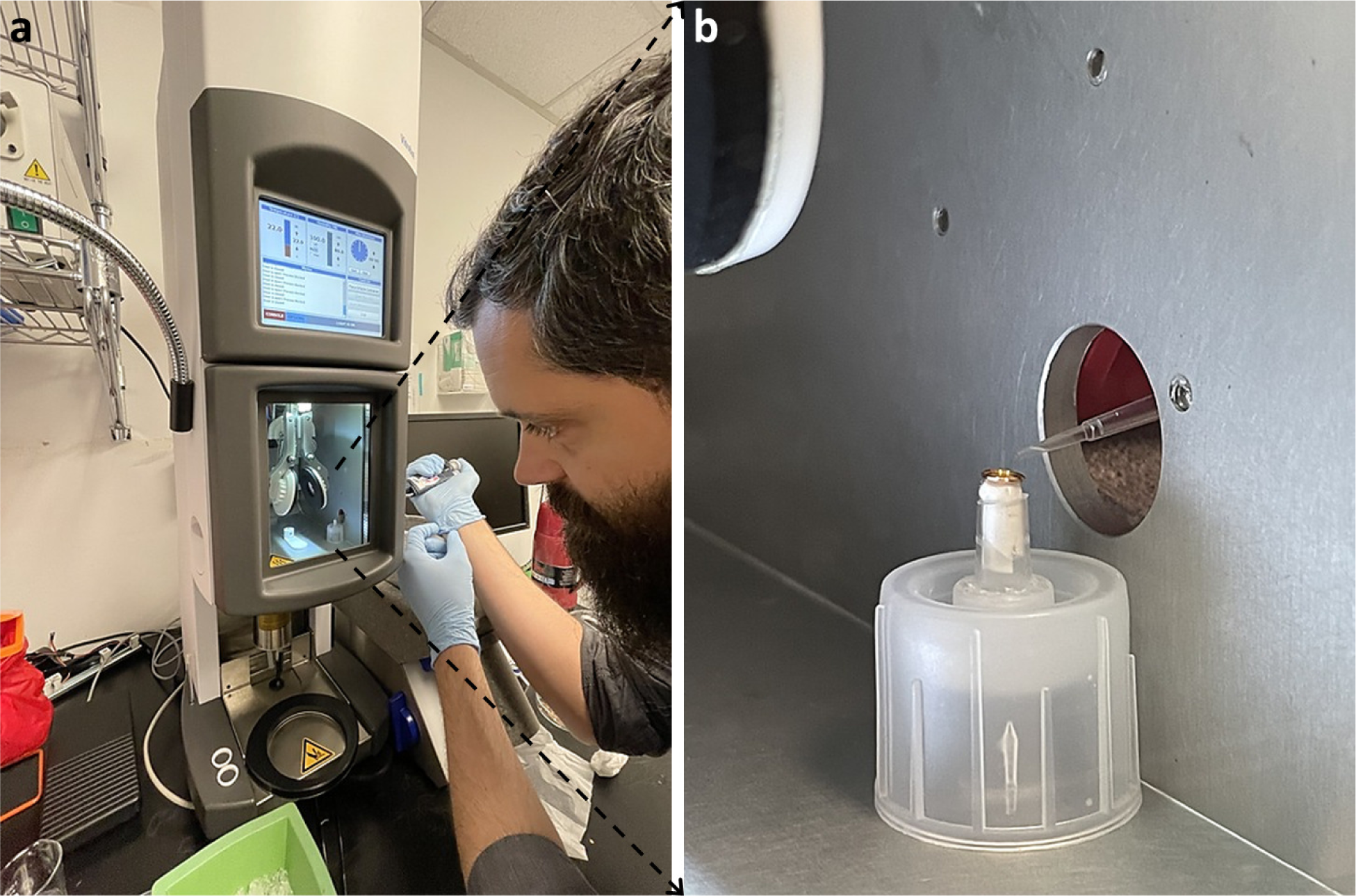
| CryoCycle-gravity development. Ongoing development of a CryoCycle setup that attempts to use gravitational force in place of manual blot force to make the method more universally reproducible. **(a)** Photo of Viacheslav Serbynovskyi testing a CryoCycle-gravity setup in a TFS Vitrobot where the chamber is only being used for humidity control. Sample is applied on top of a stabilized blotting pipette tip for through-grid wicking. Seconds after application and wicking, the grid is plunged by hand into LN2-cooled liquid ethane outside of the Vitrobot (out of view in the photo). **(b)** Zoom-in of the modified pipette tip setup, which is also visible at the bottom of the custom Allen wrench storage base shown in **Supplementary Figure 13, item #19**.

**Supplementary Figure 13.**
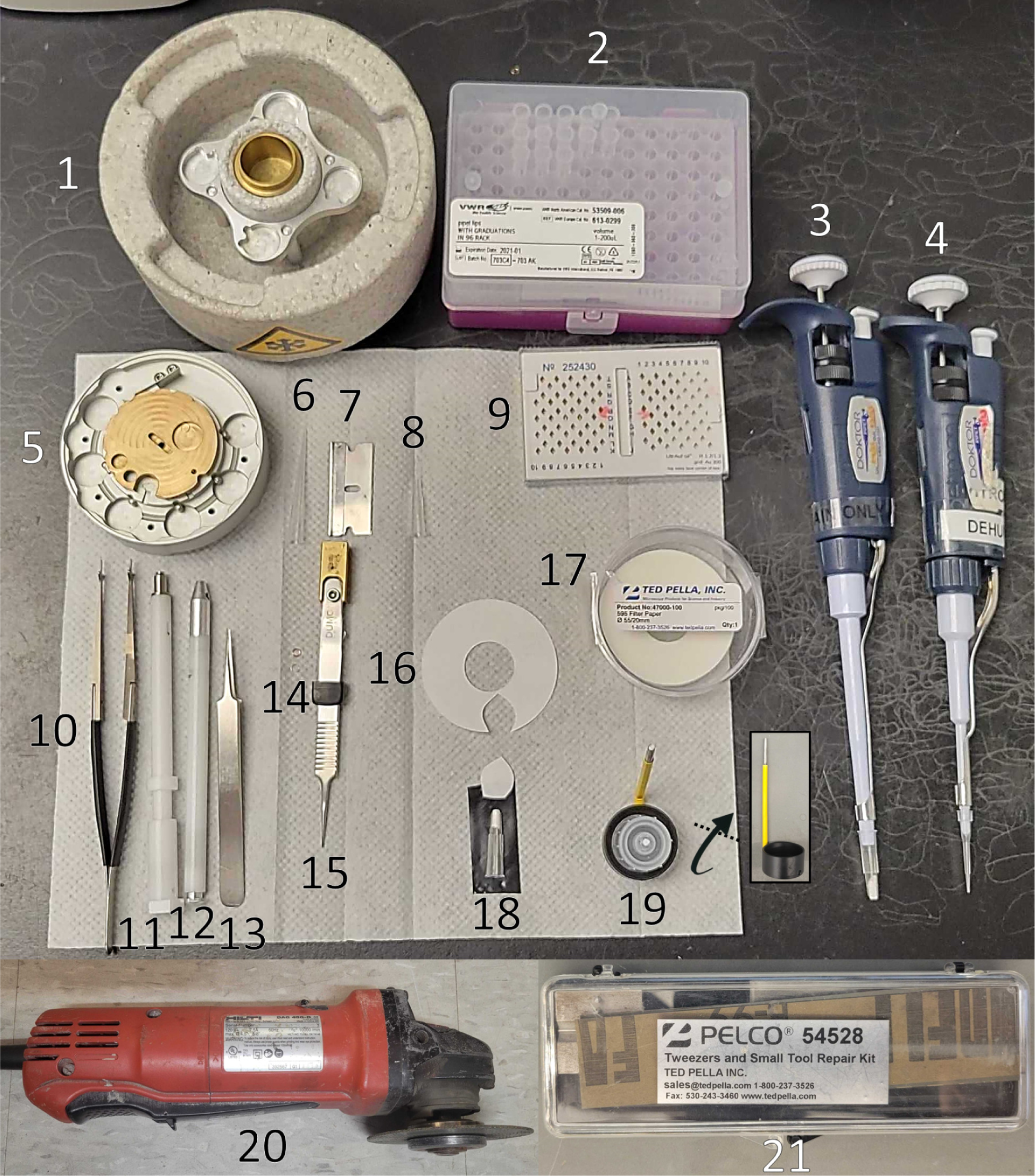
| Items recommended for preparing for CryoCycle blotting along with numbers corresponding to Supplementary Protocol 1. *Note*: CryoCycle modified pipette tips may be created with any pipette tip that has an inner diameter smaller than 3 mm; A 200 µL pipette tip is shown here. 1,000 µL and 200 µL pipette tips are shown in **Figure 1a,b**. *Note*: an autogrid clipping station (item #5) is not required; Room temperature grids can be clipped on a flat surface without a clipping station.

**Supplementary Figure 14.**
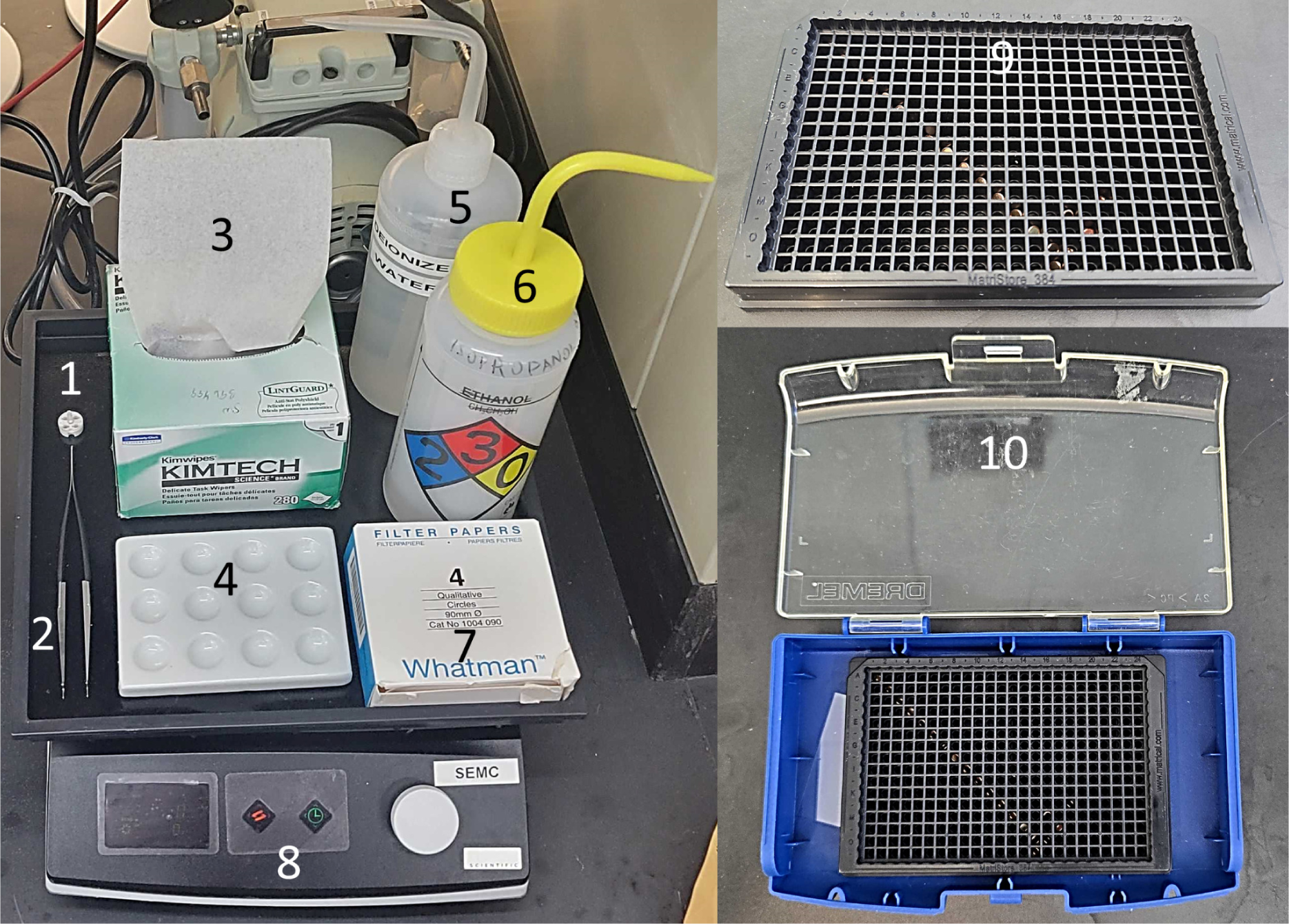
| Items recommended for the CryoCycle clipped grid washing protocol along with numbers corresponding to Supplementary Protocol 2.

**Supplementary Figure 15.**
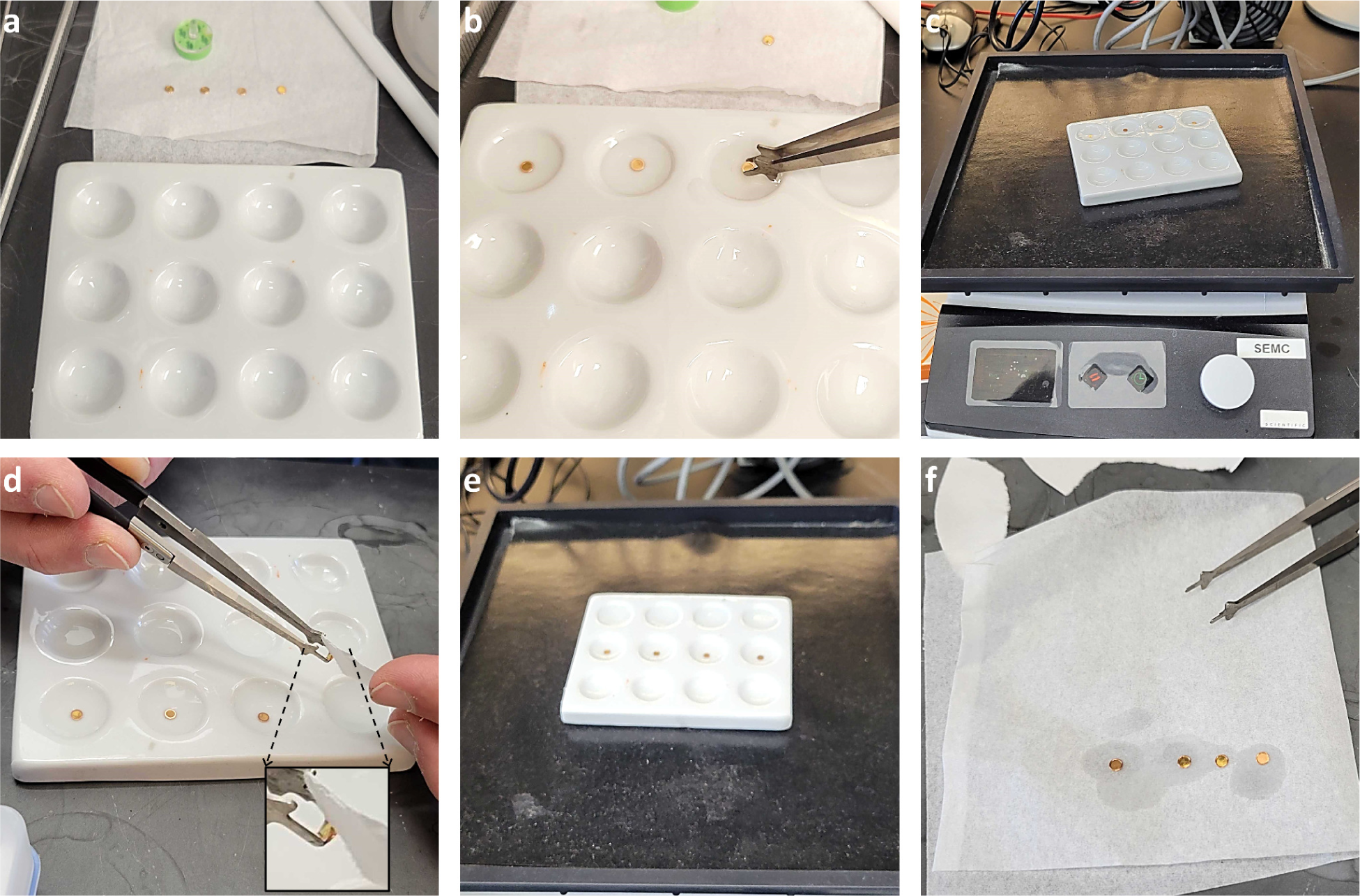
| Clipped grid washing CryoCycle protocol for reusing grids, as described in Supplementary Protocol 2. Shown here are four used clipped grids that are first warmed up **(a)**, then submerged and washed in water while shaking for 5 minutes **(b-c)**, then blotted on the side **(d)**, then submerged and washed in isopropanol while shaking for 5 minutes **(e)**, then submerged and washed in isopropanol for a second time while shaking for 5 minutes (not shown), then dried on a Kimwipe tissue **(f)**.

**Supplementary Figure 16.**
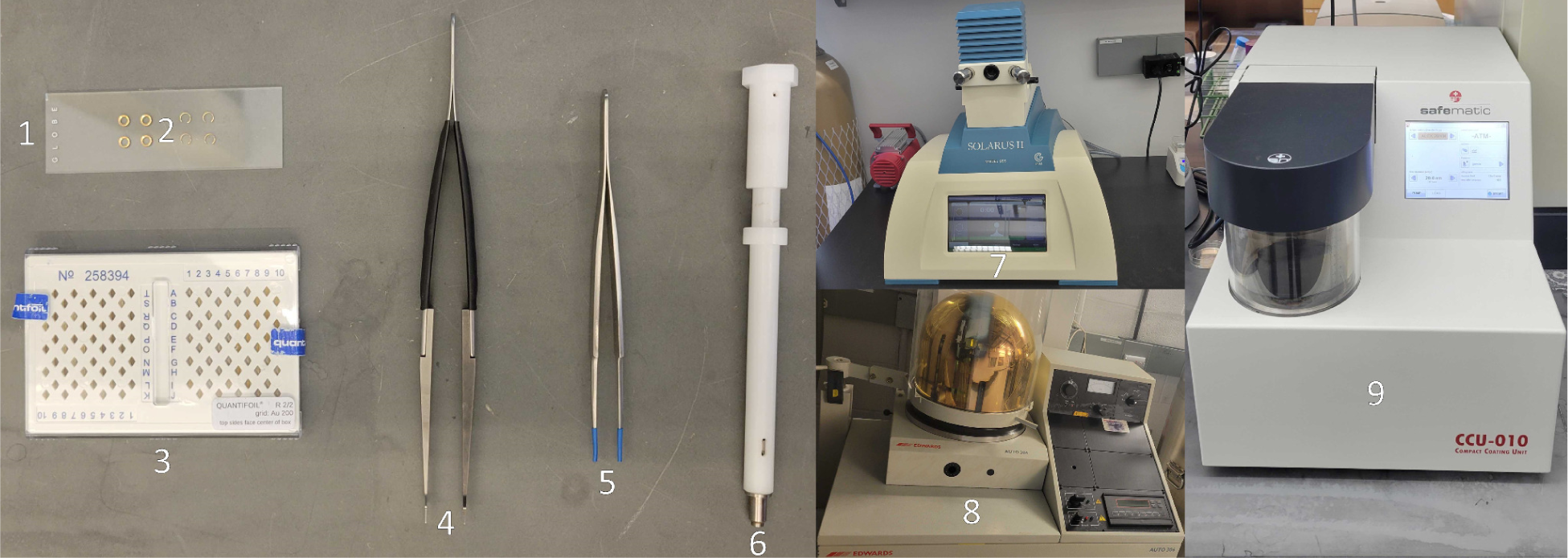
| Items recommended for preparing cells on clipped grids for use with CryoCycle cell preparation along with numbers corresponding to Supplementary Protocol 3. *Note*: The grid handling tweezers with soft tip coating (item #5) were prepared by molding heat-shrink tubing onto the tips using a flame; this addition prevents the gold-coated c-clips from being scratched by the tweezers upon insertion into the c-clip insertion tool (item #6).

## Supplementary Video 1

https://nysbc-my.sharepoint.com/:v:/g/personal/anoble_nysbc_org/Ecx-2ramF6FAkK9W5I0Vjk4BRcCW8ZXhQq_hBjEpSET9Zw?nav=eyJyZWZlcnJhbEluZm8iOnsicmVmZXJyYWxBcHAiOiJPbmVEcml2ZUZvckJ1c2luZXNzIiwicmVmZXJyYWxBcHBQbGF0Zm9ybSI6IldlYiIsInJlZmVycmFsTW9kZSI6InZpZXciLCJyZWZlcnJhbFZpZXciOiJNeUZpbGVzTGlua0NvcHkifX0&e=AXqRei

**Supplementary Video 1 | Video of Viacheslav Serbynovskyi using the CryoCycle blotting method to vitrify a single particle sample using a 1,000 μL blotting pipette tip. (0:00)** Prepare trimmed tweezers gripping a clipped grid and mounted into a plunge freezing device, LN2-cooled liquid ethane, a blotting pipette tip. **(0:19)** Pipette 3 μL of sample onto the center of the c-clip side of the clipped grid. **(0:50)** Blot the clipped grid just inside of the autogrid assembly using a blotting pipette so that the extruded filter paper makes complete contact with the face of the grid. **(1:07)** Plunge the clipped grid into liquid ethane. **(1:26)** Detach the tweezers from the plunging device, transfer the grid to the LN2, and place the grid into a clipped grid box.

## Supplementary Video 2

https://nysbc-my.sharepoint.com/:v:/g/personal/anoble_nysbc_org/ER8ZXaFLYnVAj1no8NHZ7pUB-ZW-rY2EEi7qQdLjz-x78w?nav=eyJyZWZlcnJhbEluZm8iOnsicmVmZXJyYWxBcHAiOiJPbmVEcml2ZUZvckJ1c2luZXNzIiwicmVmZXJyYWxBcHBQbGF0Zm9ybSI6IldlYiIsInJlZmVycmFsTW9kZSI6InZpZXciLCJyZWZlcnJhbFZpZXciOiJNeUZpbGVzTGlua0NvcHkifX0&e=bgAC5N

**Supplementary Video 2 | Video of Viacheslav Serbynovskyi assembling a 200 μL blotting pipette tip. (0:00)** Trim the end of a pipette tip. **(0:20)** Cut a ∼1 cm circle out from the filter paper, **(0:30)** mold it to the end of the metal rod, and **(0:38)** insert it slowly and without twisting into the large end of the trimmed pipette tip until it extends 1 mm beyond the small end. **(0:49)** Tilt the assembly about 60° to the surface of a flat filter paper and roll the edge of the extruded filter paper until it is well-rounded. **(0:54)** With the assembly tilted 90° to the surface, firmly press down. **(0:59)** Twist the rod about one eighth of a turn in one direction, then twist in the other direction while pulling the rod out of the pipette tip.

## Supplementary Protocol 1: Vitrifying Clipped Grids

This protocol is for preparing clipped grids (blotting and vitrifying) with single particle-like samples.

### Reagents and Materials (Supplementary Figure 13)

1. Vitrification dewar (e.g. Thermo Fisher Scientific, catalog number: FEI0815NB)
2. Pipette tips (e.g. VWR, catalog number: 53509-006)
3. Pipette for blotting (e.g. Rainin Classic PR-200, catalog number: 17008652)
4. Pipette for sample application (e.g. Rainin Classic PR-10, catalog number: 17008649)
5. (Optional) AutoGrid assembly workstation clipping hub (e.g. Thermo Scientific, catalog number: 1000068)
6. An example pipette tip
7. Razor (e.g. EMS, catalog number: 71960)
8. An example trimmed pipette tip
9. CryoEM grids (e.g. Quantifoil UltrAuFoil R 1.2/1.3 Au 300 mesh grids, catalog number: N1-A14nAu30-01)
10. (Optional) AutoGrid tweezers (e.g. Thermo Scientific, catalog number: 9432 909 97631)
11. C-clip insertion tool (e.g. Thermo Scientific, catalog number: 9432 909 97571)
12. Grid container tool (e.g. Thermo Scientific, catalog number: 9432 909 97671)
13. Grid handling tweezers (e.g. Dumont, catalog number: 0208-5-PO)
14. Autogrid ring and c-clip (e.g. Thermo Scientific, catalog numbers: 1036173 and 1036171)
15. Modified grid plunging tweezers (e.g. Ted Pella, catalog number: 47000-500)
16. An example filter paper with a cutout removed (e.g. Ted Pella 595 filter paper, catalog number: 47000-100)
17. Filter paper (e.g. Ted Pella 595 filter paper, catalog number: 47000-100)
18. An example assembled trimmed pipette tip with filter paper. An additional filter paper cutout is placed above the pipette tip.
19. A cylindrical metal rod (∼2 mm diameter) used for inserting the filter paper cutout into the modified pipette tip (shown in **Supplementary Figure 13** attached to a custom storage base made from an LN2 tank cap and a modified lab squeeze bottle; A modified Allen wrench that was rounded by a rotary tool is shown in the top-down view and inset side view).
20. Rotary tool to prepare tweezers, if necessary (e.g. Hilti angle grinder, catalog number: DAG 450-D)
21. Tweezer sharpening tool to prepare tweezers, if necessary (e.g. Pelco Tweezers and Small Tool Repair Kit, catalog number: 54528)
22. Several additional reagents and materials not shown in **Supplementary Figure 13** are required, including: A plunge freezing device with humidity-controlled chamber, a glow discharger, liquid nitrogen (LN2), ethane gas (99.9+% purity), cryo dewars, cryoEM sample, clipped grid boxes, clock for measuring blotting time, scissors, lab gloves, and eye protection.

### Preparation of tweezers

1. Before beginning, check if your tweezers require trimming by gripping a clipped grid on the rim as in **Supplementary Figure 6**. If the tips of the tweezers flex causing the clipped grid to rotate as in **Supplementary Figure 6b, third row**, then the tweezers require trimming. If the tips of the tweezers do not flex and the clipped grid does not rotate significantly as in **Supplementary Figure 6c, third row**, then the tweezers do not require trimming. If the tweezers require trimming, continue with the following steps.
2. Mark 2.5 mm from the tweezer tip with a permanent marker.
3. Using all proper safety measures, position the tips of the tweezers 90° to the plane of the spinning rotary tool disk. Carefully move the tweezer tips into the rotary tool disk until the tips are shaved down to the marker line.
4. Sharpen the tweezers with a sharpening tool. Ensure that the tips of the tweezers that will grip the autogrid remain square, as shown in **Supplementary Figure 7c,d**. The modified tweezers should grip a clipped grid as shown in **Supplementary Figure 6c, third row**.

### Preparation of blotting pipette tip (Supplementary Video 2)

1. Put on lab gloves. Clean all handling tools for grids and filter paper before use.
2. Trim the end of a pipette tip with a razor so that the inner diameter of the opening is 3 mm. Ensure that the end of the pipette tip is smooth.
3. Cut a ∼1 cm circle out from the filter paper.
4. Mold the filter paper cutout to the end of the rod.
5. With the filter paper cutout molded to the end of the rod, insert the assembly into the large end of the trimmed pipette tip until the filter paper extends 1 mm beyond the small end. Insert the assembly slowly and without twisting so as to not tear the filter paper.
6. Place a piece of filter paper on a flat surface. Tilt the rod, filter paper, and pipette tip assembly about 60° to the surface and roll the edge of the extruded filter paper on the flat filter paper while pushing on the rod so that the extruded filter paper is well-rounded (**Supplementary Fig. 8; Supplementary Video 2**). Tilt the assembly to be perpendicular to the flat surface and firmly press down on the rod so that the extruded filter paper is flat on the end.
7. To remove the rod, twist the rod about one eighth of a turn in one direction, then twist in the other direction while pulling the rod out of the pipette tip.
8. Store the blotting pipette tip in a clean, dry environment until ready for use. *Note*: Prepare one blotting pipette tip per grid. Do not reuse blotting pipette tips.

### Procedure for blotting and vitrifying clipped grids (Supplementary Video 1)

1. Prepare the following: trimmed tweezers, freshly glow-discharged clipped grids (grid bar side opposite to the c-clip side as shown in **Supplementary Figs. 2, 3, & 6c**), one blotting tip per clipped grid, sample for application (3 μL per grid), LN2-cooled liquid ethane in the vitrification dewar, a plunge freezing device (disable blotting, set humidity to 80+%), and a clock for measuring blotting time (a clock with audible ticks every second is recommended).
2. Grip a clipped grid with the trimmed tweezers and mount the tweezers onto the plunging device.
3. Pipette 3 μL of sample onto the center of the c-clip side of the clipped grid.
4. Use a blotting pipette tip to blot the grid inside of the autogrid ring and c-clip on the same side that the sample was applied (c-clip side) for a desired amount of time (typically 1 - 4 seconds) (**Fig. 1b**). *Note*: Ensure that the flat blotting paper fully contacts the flat grid. Ideally, the blotting pipette tip should be oriented exactly 90° relative to the plane of the grid. *Note*: Blot force is determined by the pressure on the grid exerted by the user’s hand. *Note*: Avoid contacting the blotting paper with any object other than the grid when inserting into the preparation chamber so as to not deform the flat, rounded blotting paper.
5. Plunge the clipped grid into liquid ethane.
6. Detach the tweezers from the plunging device, transfer the grid to the LN2, and place the grid into a clipped grid box. *Note*: When transferring the clipped grid from the liquid ethane to the LN2, orient the face of the clipped grid to be perpendicular to the direction of motion so as to not transfer ethane along with the clipped grid, as shown in **Supplementary Video 1**.

## Supplementary Protocol 2: Washing Clipped Grids

This protocol is for cleaning clipped grids for reuse with different single particle samples without clipped grid disassembly.

*Note*: Ensure that the structural integrity of each autogrid ring and c-clip has not been impaired between the retrieval from an autoloading microscope and insertion into the subsequent autoloading microscope so as to not damage the microscopes.

*Note*: This protocol has not been tested with cell samples.

### Reagents and Materials (Supplementary Figure 14)

1. Used clipped grids
2. Tweezers for handling autogrids (e.g. Thermo Scientific, catalog number: 9432 909 97631)
3. Low-lint tissues (e.g., Kimtech, catalog number: 34155)
4. Porcelain spot plate (e.g. Science Lab Supplies, catalog number: 3765-2)
5. Highly purified water (e.g. from a Hydro Picotap, catalog number: JKLCN0202N2H-FC)
6. Isopropanol 99+% (e.g. Sigma Aldrich, catalog number: PX1830-4)
7. Filter papers (e.g., Whatman, catalog number: 1004090)
8. Rotator/Shaker (e.g. Thermo Scientific, catalog number: 88880025)
9. (Optional) Reused grid organization tray with numbers and letters (e.g. Fluotics Matrical Matristore Microtube Rack, catalog number: MatriStore 384 Tube Rack)
10. (Optional) Storage box for the reused grid organization tray (e.g. Dremel, catalog number: 2610923299)

### Procedure for cleaning clipped grids for reuse

1. Place the used clipped grids on a tissue and let them warm up to room temperature (**Supplementary Fig. 15a**).
2. Fill a row of wells in the porcelain spot plate with purified water; one for each grid to be washed.
3. Grip each clipped grid, one by one, by the outer rim of the clip ring. Submerge each clipped grid under water in their separate wells, as shown in **Supplementary Figure 15b**.
4. Place the spot plate on the rotator and rotate at 60 rpm for 5 minutes (**Supplementary Fig. 15c**).
5. Fill the next row of wells in the spot plate with isopropanol; one for each grid being washed.
6. Grip each submerged clipped grid, one by one, by the outer rim of the clip ring. Remove each grid from the water and blot with filter paper on the side of the clip ring (i.e. not touching the grid) as shown in **Supplementary Figure 15d**. Submerge each clipped grid under isopropanol into separate wells.
7. Place the spot plate on the rotator and rotate at 60 rpm for 5 minutes (**Supplementary Fig. 15e**).
8. Repeat steps 5 through 7.
9. Place grids on a tissue to dry (**Supplementary Fig. 15f**).
10. Proceed with the Procedure in **Supplementary Protocol 1** or store the grids in an organization tray and box (**Supplementary Figure 14, items #9 & 10**) for future use.

### Recommendations

We recommend that new CryoCycle users verify the single particle CryoCycle protocol (**Supplementary Protocols 1 & 2**) in their hands as follows:

1. Prepare four new pre-clipped cryoEM grids with a standard sample, such as apoferritin (**Supplementary Protocol 1**).
2. Wash the grids for reuse (**Supplementary Protocol 2**).
3. Apply buffer as the next sample to two of the reused grids as a negative control to ensure no protein remains and apply a second standard sample, such as aldolase, as a positive control to the other two reused grids to ensure that only the second sample remains (**Supplementary Protocol 1**).
4. (Optional) Test how many times the grids can be reused (repeat steps 2 & 3).

We recommend keeping track of the following when reusing clipped grids:

- Grid type (brand, substrate, mesh size, hole size)
- Number of times reused
- Previous sample(s)
- Previous instruments & procedures used (washing protocol, plasma cleaning/glow discharge, freezing protocol, microscopes)
- Previous person(s) who handled the grid

We recommend separating clipped grids for reuse immediately after unloading from the microscope and storing them in an organization tray and box (e.g. **Supplementary Figure 14, items #9 & 10**). Use the organization tray numbering system to record the reused grid details. Washed grids can be stored in a separate organization tray.

We recommend reusing clipped grids for screening visually distinct samples to avoid misinterpretation if washing was incomplete, and only with samples that do not interact with one another.

## Supplementary Protocol 3: Preparing Pre-clipped Grids for Cells

This protocol is for preparing cell-compatible pre-clipped grids. The goal of this protocol is to gold-coat autogrid rings and c-clips so that cell samples can be grown on or applied to the pre-clipped grids without cytotoxicity from the autogrid assembly. We initially found that multiple coats of gold are required to coat autogrid rings and c-clips before clipped grid assembly, and so developed a less costly version of the protocol with an initial coating of carbon and final coating of gold.

### Reagents and Materials (Supplementary Figure 16)

1. Glass slide (e.g. EMS, catalog number: 71883-01)
2. Autogrid ring and c-clip (e.g. Thermo Scientific, catalog numbers: 1036173 and 1036171)
3. CryoEM grids (e.g. Quantifoil UltrAuFoil R 1.2/1.3 Au 300 mesh grids, catalog number: N1-A14nAu30-01)
4. (Optional) AutoGrid tweezers (e.g. Thermo Scientific, catalog number: 9432 909 97631)
5. Grid handling tweezers with soft tip coating (e.g. Dumont, catalog number: 0208-5-PO)
6. C-clip insertion tool (e.g. Thermo Scientific, catalog number: 9432 909 97571)
7. Plasma cleaner (e.g. Gatan Inc., Gatan Solarus II Model 955)
8. Gold evaporator (e.g. Edwards, Auto 306 Vacuum Coater operated at 3 kV, 50 mA)
9. Carbon evaporator (e.g. Safematic, catalog number: CCU-010)

### Procedure for preparing cell-compatible pre-clipped grids

1. Plasma clean both sides of the autogrid rings, autogrid c-clips, and grids (oxygen and argon for 7 seconds).
2. Place the autogrid rings and c-clips in the carbon evaporator and coat them with ∼20 nm of carbon on both sides.
3. Place the autogrid rings and c-clips in the gold evaporator and coat them with several nanometers of gold on both sides.
4. Clip grids using the carbon+gold coated autogrid rings and c-clips at room temperature. Use the soft tip coated tweezers for inserting the c-clip into the c-clip insertion tool.
5. Plasma clean both sides of the pre-clipped grids (oxygen and argon for 7 seconds).
6. Grow or apply cells to the grid bar side of the grid (opposite to the c-clip side as shown in **Supplementary Fig. 3**).
7. Proceed with the Procedure in **Supplementary Protocol 1**, except skip step 3 and blot from the side opposite to sample application as shown in **Supplementary Figure 3**.

